# Soil Protists in Three Neotropical Rainforests are Hyperdiverse and Dominated by Parasites

**DOI:** 10.1101/050997

**Authors:** Frédéric Mahé, Colomban de Vargas, David Bass, Lucas Czech, Alexandros Stamatakis, Enrique Lara, David Singer, Jordan Mayor, John Bunge, Sarah Sernaker, Tobias Siemensmeyer, Isabelle Trautmann, Sarah Romac, Cédric Berney, Alexey Kozlov, Edward A.D. Mitchell, Christophe V. W. Seppey, Elianne Egge, Guillaume Lentendu, Rainer Wirth, Gabriel Trueba, Micah Dunthorn

**Affiliations:** Department of Ecology, University of Kaiserslautern, Erwin-Schrödinger-Straße, 67663 Kaiserslautern, Germany; CNRS, UMR 7144, Station Biologique de Roscoff, Place Georges Teissier, 29680 Roscoff, France; Sorbonne Universités, UPMC Univ Paris 06, UMR 7144, Station Biologique de Roscoff, Place Georges Teissier, 29680 Roscoff, France; Department of Life Sciences, The Natural History Museum London, Cromwell Road, London SW7 5BD, United Kingdom; Centre for Environment, Fisheries & Aquaculture Science (Cefas), Barrack Road, The Nothe, Weymouth, Dorset DT4 8UB, United Kingdom; Heidelberg Institute for Theoretical Studies, Schloß-Wolfsbrunnenweg, 69118 Heidelberg, Germany; Institute for Theoretical Informatics, Karlsruhe Institute of Technology, Am Fasanengarten, 76128 Karlsruhe, Germany; Laboratory of Soil Biodiversity, Université de Neuchâtel, Rue Emile-Argand, 2000 Neuchâtel, Switzerland; Department of Forest Ecology and Management, Swedish University of Agricultural Sciences, Skogsmarksgränd, 90183 Umeå, Sweden; Department of Computer Science, Cornell University, Gates Hall, Ithaca, New York 14853, USA; Department of Statistical Science, Cornell University, Malott Hall, Ithaca, New York 14853, USA; Jardin Botanique de Neuchâtel, Pertuis-du-Sault, 2000 Neuchâtel, Switzerland; Department of Biosciences, University of Oslo, Blindernveien, 0316 Oslo, Norway; Department of Plant Ecology and Systematics, University of Kaiserslautern, Erwin-Schrödinger-Straße, 67663 Kaiserslautern, Germany; Instituto de Microbiología, Universidad San Francisco de Quito, Diego de Robles, Quito, Ecuador

## Abstract

Animal and plant richness in tropical rainforests has long intrigued naturalist. More recent work has revealed that parasites contribute to high tropical tree diversity (Bagchi *et al.*, 2014; Terborgh, 2012) and that arthropods are the most diverse eukaryotes in these forests (Erwin, 1982; Basset *et al.*, 2012). It is unknown if similar patterns are reflected at the microbial scale with unicellular eukaryotes or protists. Here we show, using environmental metabarcoding and a novel phylogeny-aware cleaning step, that protists inhabiting Neotropical rainforest soils are hyperdiverse and dominated by the parasitic Apicomplexa, which infect arthropods and other animals. These host-specific protist parasites potentially contribute to the high animal diversity in the forests by reducing population growth in a density-dependent manner. By contrast, we found too few Oomycota to broadly drive high tropical tree diversity in a host-specific manner under the Janzen-Connell model (Janzen, 1970; Connell, 1970). Extremely high OTU diversity and high heterogeneity between samples within the same forests suggest that protists, not arthropods, are the most diverse eukaryotes in tropical rainforests. Our data show that microbes play a large role in tropical terrestrial ecosystems long viewed as being dominated by macro-organisms.

---

Since the works of early naturalists such as von Humboldt and Bonpland (von Humboldt and Bonpland, 1852), we have known that animals and plants in tropical rainforests are exceedingly species rich. For example, one hectare can contain more than 400 tree species (Valencia *et al.*, 1994) and one tree can harbour more than 40 ant species (Wilson, 1987). This hyperdiversity of trees has been partially explained by the Janzen-Connell model (Janzen, 1970; Connell, 1970), which hypothesizes that host-specific predators and parasites reduce plant population growth in a density-dependent manner (Bagchi *et al.*, 2014; Terborgh, 2012). Sampling up in the tree canopies and below on the ground has further found that arthropods are the most diverse eukaryotes in tropical rainforests (Erwin, 1982; Basset *et al.*, 2012). The focus on eukaryotic macro-organisms in these studies is primarily because they are familiar and readily observable to us. We do not know if the less familiar and less readily observable protists—microbial eukaryotes excluding animals, plants, and fungi (Pawlowski *et al.*, 2012)—inhabiting these same ecosystems exhibit similar diversity patterns.

To evaluate if macro-organismic hyperdiversity patterns are reflected at the microbial scale, we sequenced 132.3 million cleaned V4 reads from soils sampled in Costa Rica, Panama, and Ecuador. Of the 50.1 million reads assigned to the protists, 75.3% had a maximum similarity of <80% to references in the Protist Ribosomal Reference (PR2) database (Guillou *et al.*, 2013) (Fig. 1, Extended Data Fig. 1). By contrast, only 3.1% were <80% similar to the PR2 database in our re-analysis of 367.8 million V9 reads from the open oceans, and most of the V4 and V9 reads from other marine environments were likewise highly similar to the PR2 database (Fig. 1, Extended Data Fig. 1). Reads <80% similar to known references are usually considered spurious and removed in environmental protist sequencing studies (Stoeck *et al.*, 2010). Three quarters of our rainforest soil protist data would be discarded if we applied this conservative cleaning step. However, PR2 and similar databases are biased towards marine and temperate terrestrial reference sequences. To solve this problem, reads were dereplicated into 10.6 million amplicons (i.e. strictly identical reads to which an abundance value can be attached), then placed onto a phylogenetic tree inferred from 512 full-length reference sequences from all major eukaryotic clades (Fig. 2, Extended Data Fig. 2). We conservatively retained operational taxonomic units (OTUs) whose most abundant amplicon fell only within known clades with a high likelihood score. This novel phylogeny-aware cleaning step effectively discarded incorrect, too short or highly divergent amplicons (Dunthorn *et al.*, 2014), resulting in the removal of only 6.8% of the reads and 7.7% of the OTUs.

**Fig. 1.**
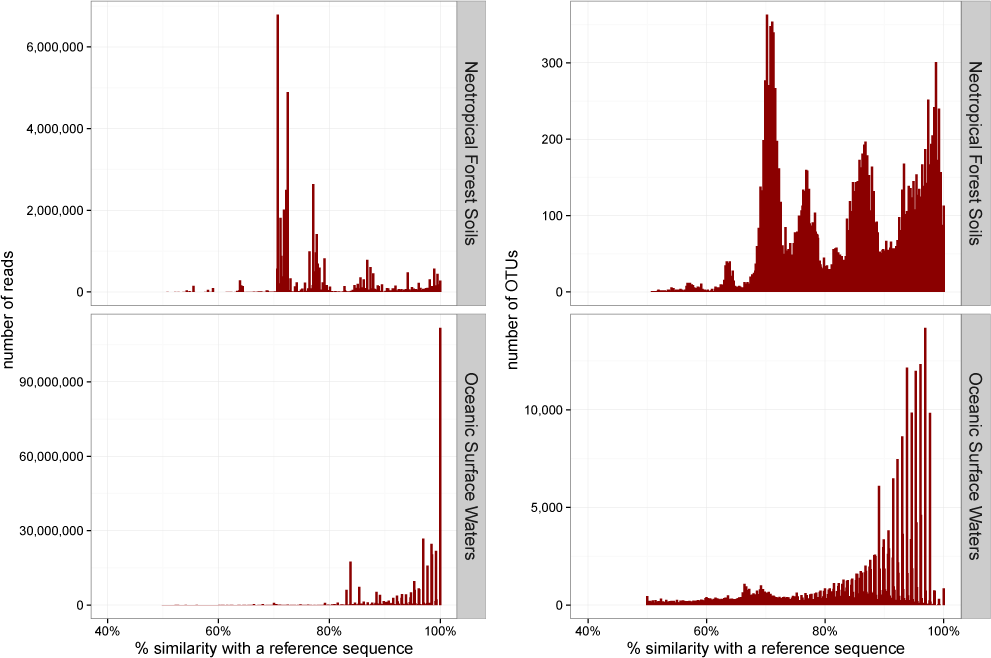
Similarity of protists to the taxonomic reference database PR2 (Guillou *et al*., 2013). In contrast to marine data, most of the reads and OTUs from the Neotropical rainforest soils were <80% to references in the PR2 database. Only 8.1% soil reads had a similarity *≥*95% with references, while 68.1% of the marine reads from the Tara-Oceans open oceans had a similarity *≥*95% with reference sequences.

**Fig. 2.**
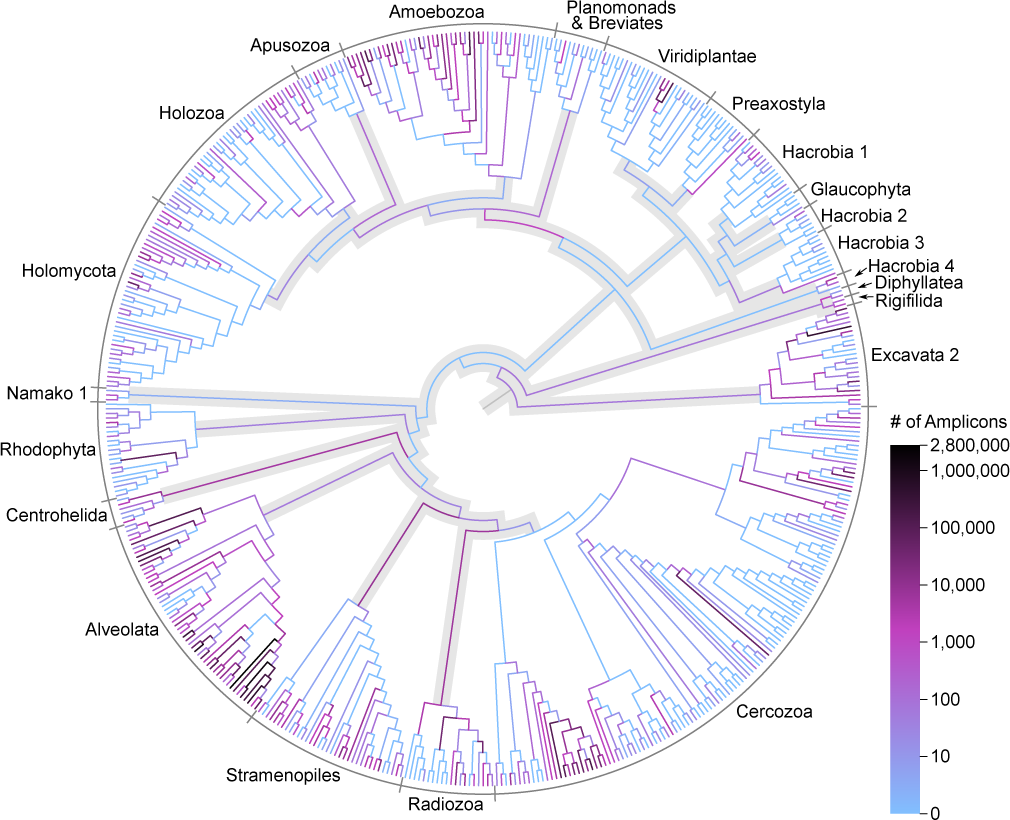
Phylogenetic placement of Neotropical soil protist reads on a taxonomically unconstrained global eukaryotic tree. Reads were dereplicated into strictly identical amplicons. Inferred relationships between these major taxa may differ from those obtained with phylogenomic data. Alveolate includes Apicomplexa and Ciliophora; Holozoa includes animals; Holomycota includes fungi. Branches and nodes outside of known clades are shaded in gray. In our conservative approach, OTUs placed along shaded branches were discarded.

The remaining protist reads from the rainforest soils clustered into 26,860 OTUs. As in the marine plankton (de Vargas *et al.*, 2015), more protist OTUs were detected than animals (4,374, of which 39% were assigned to the Arthropoda), plants (3,089), and fungi (17,849) combined (Extended Data Table 1a). Some of these hyperdiverse protists and other eukaryotes found in the soils could be a shadow of the tree-canopy communities from cells that have rained down from above. Taxonomic assignment of the protists showed that 84.4% of the reads and 50.6% of the OTUs were affiliated to the Apicomplexa (Fig. 3). Apicomplexa are widespread parasites of animals (Desportes and Schrével, 2013; Perkins *et al.*, 2009), but their close relatives are free-living (Janouškovec *et al.*, 2015). Apicomplexa read- and OTU-abundances were much lower in marine and other terrestrial environments (Extended Data Fig. 3a). A high abundance of Apicomplexa was also observed in rainforest soil samples amplified with primers designed to target just the closely related Ciliophora (Extended Data Fig. 3b).

**Fig. 3.**
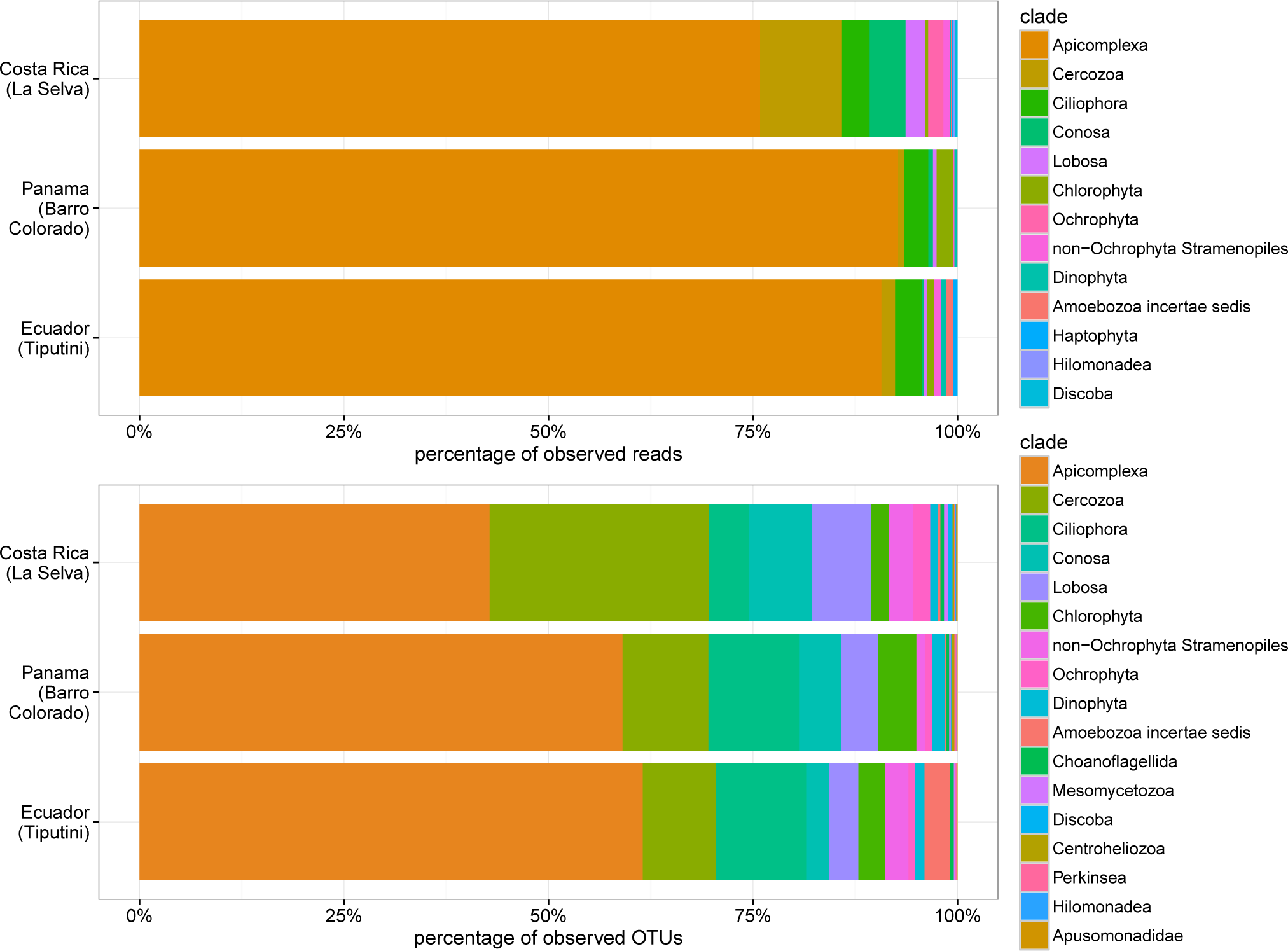
Taxonomic identity and relative abundances of soil protist reads and OTUs in three Neotropical rainforests. Taxa shown represent *>*0.1% of the data. Taxonomy is at the 3rd level of the PR2 database. Between 76.6% and 93.0% of the reads, and between 43.1% and 61.5% of the OTUs, per forest were assign to the Apicomplexa (in orange).

We placed Apicomplexa OTUs into a more focused phylogeny inferred from 190 full-length reference sequences from all major Alveolate clades. While the OTUs were generally distributed across the whole tree (Extended Data Fig. 4), 80.2% of them grouped with the gregarines, which predominantly infect arthropods and other invertebrates (Desportes and Schrével, 2013). About 23.8% of these gregarine OTUs placed within the lineage formed by the millipede parasite Stenophora, and the insect parasites Amoebogregarina, Gregarina, Leidyana, and Protomagalhaensia. Many other OTUs placed within gregarine lineages thought to be primarily parasites of marine annelids and polychaetes. Non-gregarine Apicomplexa OTUs largely grouped with the blood parasites Plasmodium (which cause malaria) and close relatives, some of which cycle through arthropods and vertebrates such as birds.

These dominating protist parasites potentially contribute to high animal diversity in the rainforests like other parasites contribute to high tree diversity in the Janzen-Connell model (Janzen, 1970; Connell, 1970). The usually host-specific Apicomplexa, although switching is known (Desportes and Schrével, 2013; Perkins *et al.*, 2009; Ellis *et al.*, 2015; Rueckert *et al.*, 2011), infect animals (e.g., Asghar *et al.* 2015; Bouwma *et al.* 2005) and may limit population growth of species that become locally abundant. In our tentative model, we expect Apicomplexa to dominate both the below- and above-ground protist communities in tropical forests worldwide, most animal populations in those forests will have apicomplexan parasites, and animals that become locally dominant will have escaped from their apicomplexan enemies. This relationship between animals and apicomplexans could also be reciprocal, with each group contributing to the diversity of the other.

The second through fifth most diverse protist were the predominantly predatory Cercozoa, Ciliophora, Conosa and Lobosa (Fig. 3). Photo- or mixo-trophic Chlorophyta, Dinophyta, Haptophyta, and Rhodophyta were also detected. Haptophyta are mostly marine, for example, but phylogenetic placement of these OTUs showed most had close relationships to freshwater species (Extended Data Fig. 5). Only 131 OTUs were assigned to the Oomycota. These efficient parasites, which have ciliated stages that disperse through soils, have long been thought to drive the Janzen-Connell model. However, their degree of host-specificity is unknown for most species (Freckleton and Lewis, 2006), and a fungicide study in Belize documented a non-significant effect by the Oomycota (Bagchi *et al.*, 2014). If Oomycota broadly drive the Janzen-Connell model, then we can expect their diversity to be very high, mirroring tree diversity, but we found too few Oomycota to be species-specific plant parasites under this model and many of them were closely related to taxa that infect animals (Extended Data Fig. 6).

We estimated less than 1,732 unobserved OTUs with each forest (Extended Data Table 1b), suggesting that the sequencing depth enabled detection of much of the total diversity within each sample. Bray-Curtis dendrograms and non-metric multidimensional scaling (Extended Data Fig. 7a,b) showed that the OTU composition was slightly more similar among the samples from Costa Rica and Panama than those from Ecuador, potentially reflecting geography. However, the Jaccard similarity index estimated high OTU heterogeneity between samples even within the same forest, and within-forest OTU rarefaction curves based on the sample accumulation further estimated logarithmic increases without plateaus (Extended Data Fig. 7c,d). Overall, this indicates that our sampling effort has unveiled only a fraction of the protist hyperdiversity in the three rainforests.

Just as there is high richness at the macro-organismic scale, the high OTU numbers and high heterogeneity among samples show that there are similar patterns of hyperdiversity with protists at the microbial scale. Given the low similarity of protist compositions between samples and the 6-fold ratio of protist to animal OTUs, a concerted and comparable count will likely show that protists are more diverse than arthropods. This would certainly be the case if, as we suggest, every arthropod species has at least one apicomplexan parasite, and the Apicomplexa are only one part of total protist hyperdiversity. If protists are the most diverse eukaryotes in tropical rainforests it would not be due to inordinate speciation in just a few clades since the mid-Phanerozoic (e.g., the beetles Farrell 1998), but because of the diversification of rich and functionally complex protist lineages beginning in the early Proterozoic (Knoll, 2014), which built up the multifaceted unseen foundation of these now familiar macroscopic terrestrial ecosystems.

## METHODS SUMMARY

We sampled soils in 279 locations in a variety of lowland Neotropical forest types in: La Selva Biological Station, Costa Rica; Barro Colorado Island, Panama; and Tiputini Biodiversity Station, Ecuador (Supplementary File 1). By amplifying DNA (Extended Data Fig. 8) extracted from the soils with general primers for the V4 region of 18S rRNA and sequencing with Illumina MiSeq, we were able to detect most eukaryotic groups and assess the diversity and the relative dominance of free-living and parasitic lineages. Reads were clustered into OTUs with Swarm (Mahé *et al.*, 2015) (Extended Data Fig. 9). To provide a comparative context, we applied the same bioinformatics pipeline to re-analyse the Tara-Oceans consortium’s data of the V9 region of 18S rRNA (de Vargas *et al.*, 2015), sequenced 40 European near-shore marine samples with V4 and V9 primers, and sequenced 20 soil samples with ciliate-specific primers. Phylogenetic placements used the Evolutionary Placement Algorithm (Berger *et al.*, 2011) (Supplementary File 2), and alpha diversity was estimated with Breakaway (Willis and Bunge, 2015). All reads were deposited at GenBank’s SRA bioproject #SUB582348.

## ACKNOWLEDGEMENT

We thank Thorsten Stoeck, Laura Katz, Håvard Kauserud, and Owen T. Lewis, and David Montagnes for helpful comments.

## Funding

Funding primarily came from the Deutsche Forschungsgemeinschaft grant DU1319/1-1 to M.D, and the Klaus Tschira Foundation to L.C., A.S and A.K. Funding also came from: Natural Environment Research Council (NERC) grants NE/H000887/1 and NE/H009426/1 to D.B., the National Science Foundation’s International Research Fellowship Program (OISE-1012703) and the Smithsonian Tropical Research Institute’s Fellowship program to J.R.M., the Swiss National Science Foundation (310003A 143960) and the Swiss Federal Office for the Environment and Swiss National Science Foundation to E.L. and E.A.D.M. The authors gratefully acknowledge the Gauss Centre for Supercomputing e.V. (www.gauss-centre.eu) for funding this project by providing computing time on the GCS Supercomputer SuperMUC at Leibniz Supercomputing Centre (LRZ, www.lrz.de).

